# Adaptation and competition in deteriorating environments

**DOI:** 10.1101/734509

**Authors:** Romana Limberger, Gregor F. Fussmann

## Abstract

Evolution might rescue populations from extinction in changing environments. Using experimental evolution with microalgae, we investigated if competition influences adaptation to an abiotic stressor, and vice versa, if adaptation to abiotic change influences competition. In a first set of experiments, we propagated monocultures of five species with and without increasing salt stress for ~180 generations. When assayed in monoculture, two of the five species showed signatures of adaptation, that is, lines with a history of salt stress had higher population growth rates at high salt than lines without prior exposure to salt. When assayed in mixtures of species, however, only one of these two species had increased population size at high salt, indicating that competition can alter how adaptation to abiotic change influences population dynamics. In a second experiment, we cultivated two species in monocultures and in pairs, with and without increasing salt. While we found no effect of competition on adaptation to salt, our experiment revealed that evolutionary responses to salt can influence competition. Specifically, one of the two species had reduced competitive ability in the no-salt environment after long-term exposure to salt stress. Collectively, our results highlight the complex interplay of adaptation to abiotic change and competitive interactions.

## Introduction

Abiotic environmental change can render populations maladapted, leading to their decline, and potentially, extinction [1–3]. Population decline may be slowed or arrested by adaptation [4, 5], that is, by phenotypic change that improves fitness and has a genetic basis [6]. Both laboratory and field-based studies have demonstrated that organisms can adapt rapidly to a variety of abiotic drivers, including acidification [7], salt stress [8], pollution [9], nutrient limitation [10], and warming [11, 12]. Experiments have also demonstrated evolutionary responses to biotic drivers, such as competition [13–15] and predation [16]. However, only few experiments have explored the interactive effects of abiotic change and biotic interactions on adaptation [16–19]. Yet, species interactions might influence (i) whether populations adapt to abiotic change [16–18], and (ii) whether adaptation increases population size and persistence [20]. Moreover, within communities, the ecological consequences of adaptation to abiotic change could go beyond effects on population size, and include effects on species interactions and community dynamics [21]. This potentially complex interplay of biotic interactions and adaptation to abiotic change limits our understanding of whether evolution can contribute to the maintenance of diversity in changing environments.

Theory suggests several mechanisms whereby interspecific competition can influence adaptation to deteriorating environmental conditions. First, competition can constrain adaptation to environmental change by reducing population size, and thus the amount of standing genetic variation [22] and the supply of beneficial mutations [23]. Second, competition can reduce the time available for adaptation as species are outcompeted by competitors with broader environmental tolerance [24]. Third, competition can alter the strength and direction of selection [25]; depending on whether competition and environmental change impose selection on a trait in the same or in the opposite direction, competition can speed up or hinder adaptation to environmental change [25]. Finally, negative genetic correlations between traits that mediate competition and stress tolerance could constrain adaptation in environments with both biotic and abiotic selective agents [26]. Only few experiments have analysed evolutionary responses to a factorial manipulation of abiotic change and biotic interactions to test these theoretical predictions [16–19]. Moreover, most of these empirical tests investigated biotic interactions other than interspecific competition [16, 17], or environmental amelioration rather than deterioration [17, 18, but see 19]. In our study, however, we focused on interspecific competition, a key biotic interaction, and its interplay with adaptation to deteriorating conditions, that is, to environmental changes that have a negative effect on fitness (e.g. acidification, pollution).

Competition could not only alter how species adapt to abiotic change, but could also modulate the effect of adaptation on population dynamics. When competition alters the fitness landscape generated by abiotic change, adaptation of a focal species in isolation may not translate into increased population size in a community context. Accordingly, an experiment that manipulated CO_2_ and diversity of a plant community found adaptation to elevated CO_2_ only when plants were assayed in the same community context in which selection had occurred [18]. Similarly, an experiment with a marine alga revealed that adaptation to CO_2_ enrichment in isolation was not associated with increased population size within communities [20]. The scenario of adaptation in isolation followed by competition might seem artificial, but it is a plausible scenario in a spatial context. Species could adapt to changes in their local environment before immigration of better-adapted competitors from the regional species pool [27, 28]. Such changes in competitive interactions could prevent that adaptation of a focal species to abiotic change translates into increased population size.

When investigating the ecological consequences of adaptation to an abiotic stressor, the focus is often on effects on population size and persistence, but within communities, adaptation of a focal species to the local environment may affect interactions with other species [29–31]. Evolution can influence community dynamics directly, by altering traits that underlie species interactions, and indirectly, by influencing population dynamics [32, 33]. In a scenario with an abiotic stressor as the selective agent, adaptation could influence species interactions if traits that confer adaptation to the stressor are correlated with traits that affect biotic interactions. Some traits may even simultaneously mediate stress tolerance and species interactions. For example, in *E. coli* reduced membrane permeability increases resistance to antibiotics but decreases the ability to take up nutrients, such that adaptation to antibiotics can result in lower competitive ability [34]. In addition to such trait-mediated effects, adaptation of a focal species to an abiotic stressor might influence biotic interactions by increasing its population size. Despite this range of possibilities of how adaptation to an abiotic stressor could affect biotic interactions, its ecological consequences have rarely been considered in a community context.

Here we used experimental evolution with freshwater algae to investigate if adaptation to abiotic change influences competitive interactions, and, vice versa, if competition influences adaptation to deteriorating environmental conditions. We used salt as a stressor because it has a negative effect on the fitness of freshwater algae, and because it can easily be manipulated without confounding changes in other abiotic factors. We did not intend to mimic a specific environmental change scenario observed in natural systems; rather we wanted to investigate conceptual questions about the interplay of competition and adaptation to an abiotic stressor. We conducted two sets of experiments (Fig. 1): in the first experiment we investigated if adaptation to salt in monoculture translated into increased population size (i.e. cell number) within salt-stressed mixtures of species, to test the hypothesis that competition can alter the effect of adaptation on population dynamics. In the second experiment we tested the hypotheses that interspecific competition constrains adaptation to salt stress, and that a history of salt stress leads to increased competitive ability at high salt.

**Figure 1:**
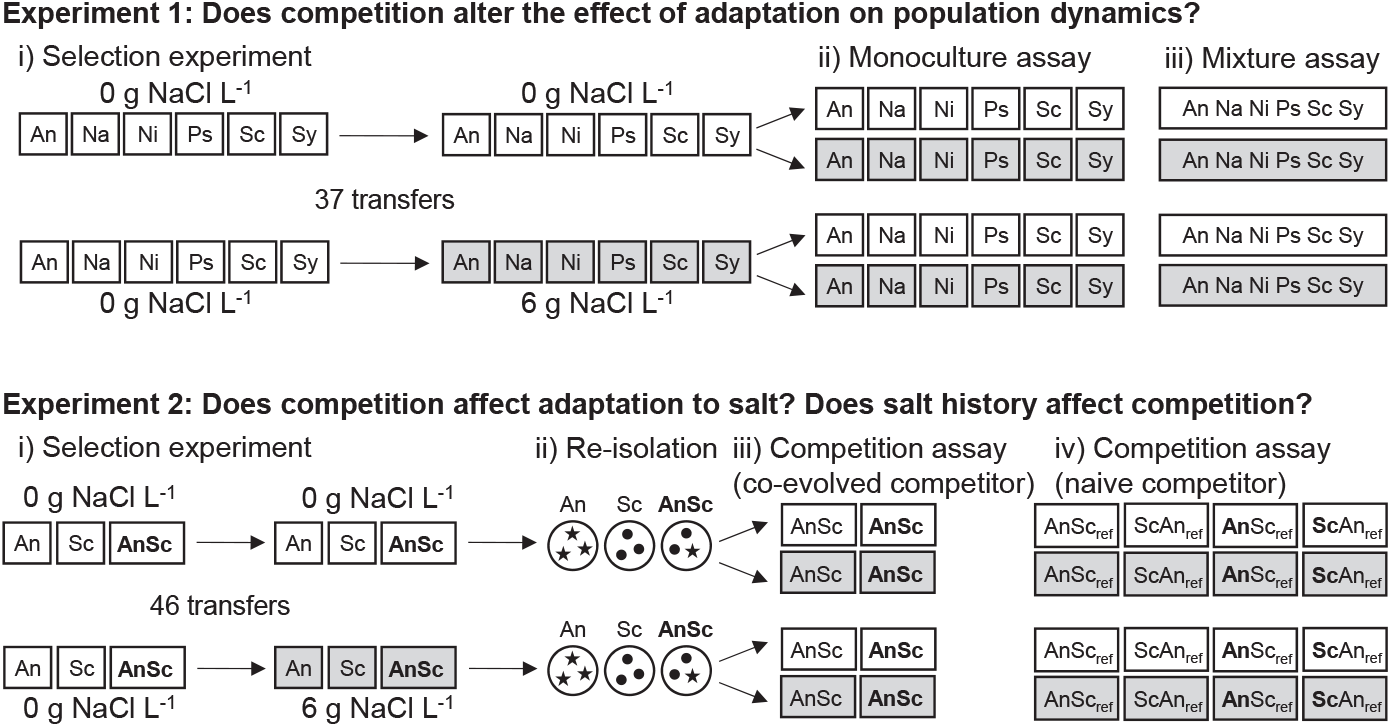
Experimental design. In Experiment 1, we propagated monocultures of six species of microalgae with or without increasing salt for ~ 185 generations, and then quantified growth rate in monoculture and population size in mixtures of species with and without salt. White boxes are flasks without salt, grey boxes are flasks with salt. In Experiment 2, we propagated two species in monoculture and in pairs, with and without salt. After ~ 230 generations we re-isolated the species on agar plates and quantified the chlorophyll-a concentration (as proxy for biomass) of all lines when competing either with a competitor of the same salt and competition history (co-evolved competitor) or with a reference line (naïve competitor; salt history: no salt, competition history: monoculture). Bold species abbreviations denote lines cultivated in pairs during the selection experiment.

## Material and methods

### Model organisms

We tested our hypotheses using six species of microalgae from three different taxonomic groups: the cyanobacteria *Synechococcus leopoliensis* (Canadian Phycological Culture Centre (CPCC) 102) and *Anabaena variabilis* (CPCC 105), the diatoms *Navicula pelliculosa* (CPCC 552) and *Nitzschia palea* (CPCC 160), and the chlorophytes *Pseudokirchneriella subcapitata* (CPCC 37) and *Scenedesmus acutus* (CPCC 10). We chose these six species because they were available as axenic (i.e. bacteria-free) cultures and because we knew from previous experiments that they grow well under the same experimental conditions [35, 36]. The algae were cultivated in Bold’s basal medium [37], which was modified by adding silicate (0.58 g L^−1^ of Na_2_SiO_3_) and vitamins [38] to allow growth of diatoms. In all experiments, the algae were cultivated at 25 °C with light continuously provided at 100 μE m^−2^ s^−1^. Cultures were axenic at the start of the experiment and propagated under sterile conditions. At the end of the experiment, we plated a subset of the cultures (18 cultures of *Anabaena* and *Scenedesmus*) on agar and observed no bacterial contaminations. We did not make cultures isogenic at the start of the experiment; the observed evolutionary responses could thus have resulted either from *de novo* mutations or from sorting of standing genetic variation.

In a preliminary experiment, we measured salt tolerance of the six species (Fig. S1), and found a low limit of salt tolerance in *Nitzschia* and *Pseudokirchneriella* (~6 g NaCl L^−1^), intermediate salt tolerance in *Anabaena, Scenedesmus* and *Navicula* (limit: ~14 g NaCl L^−1^), and high salt tolerance in *Synechococcus* (limit: > 20 g NaCl L^−1^).

### Adaptation to salt in monoculture

In a serial transfer experiment, we propagated monocultures of the six species with and without increasing concentrations of NaCl (Fig. 1, Fig. S2). The algae were cultured in 125-ml glass flasks filled with 50 ml of Bold’s medium and continuously shaken at 250 rpm. Every 3.5 days, we transferred 1 ml of culture to 50 ml of fresh medium for a total of 37 transfers (~ 185 generations; one transfer corresponding to five generations). For each of the six species, three replicate control lines were cultivated without addition of salt and three replicate selection lines were cultivated with increasing salt. We increased the salt concentration in the selection lines by 0.25 g L^−1^ every transfer until reaching 4 g NaCl L^−1^ at transfer 16 (Fig. S2). We maintained this salt concentration for 10 transfers and then continued the increase in salt until reaching the final concentration of 6 g NaCl L^−1^ at transfer 34, which we maintained for three more transfers. This salt concentration had a distinctly negative effect on the growth rates of species with lowest salt tolerance but still allowed persistence of all selection lines. A further increase in salt concentration would have probably resulted in extinction of species with low tolerance and would thus have reduced the number of species in the mixture assay.

The goal of the salt treatment was to simulate an environmental change scenario as it would be experienced by the members of a community; we thus exposed all species to the same change in environmental conditions rather than exposing each species to its respective limit of salt tolerance. Because the species varied in salt tolerance (Fig. S1), they differed in how strongly they were stressed by the salt treatment. Similarly, because the species varied in growth rates and population sizes, they differed in the potential to adapt to salt. Such species-specific differences are a property of any community and we thus made no attempt to reduce these differences (e.g. by transferring equal cell numbers rather than equal volume).

After 37 transfers (~ 185 generations), we measured population growth rate of each line in the two final salt environments (0 and 6 g NaCl L^−1^) to test if the species had adapted to salt stress. Before starting the growth assay, we cultivated all lines at identical environmental conditions for three transfer cycles, that is two cycles in the ancestral environment (0 g NaCl L^−1^) and one cycle in the assay environment (0 and 6 g NaCl L^−1^), to reduce non-evolutionary effects and pre-condition the lines to the assay conditions. We then inoculated flasks containing 50 ml medium of either 0 or 6 g NaCl L^−1^ with 1 ml of acclimated culture and measured absorbance through time. Depending on the time of individual cultures to reach carrying capacity, we measured absorbance over 2.5 to 7.5 weeks with 23 to 34 measurements. At each measurement, we removed 200 μl of culture from each flask into 96-well plates and measured absorbance at 660 nm on an optical plate reader (Synergy-HT, BioTek, Winooski, VT). As done in other studies [8, 35], we calculated population growth rates (see below) from raw absorbance values, i.e., we did not use standard curves to transform absorbance to cell density. We thus did not take into account that changes in cell size in response to salt might have influenced absorbance.

### Mixture assay

To test if a history of salt stress can influence population size of species within salt-stressed mixtures of species, we assembled mixtures either of control or of selection lines, transplanted the mixtures into the two final salt environments of the selection experiment, and quantified species abundances over four transfers. Prior to the mixture assay, the lines had been cultivated for 37 transfers with or without increasing salt, followed by two transfers in the ancestral environment (see above). We assembled mixtures with equal biovolume of the six species. To this end, we determined species abundances in the monocultures by counting Lugol-fixed samples with an inverted microscope, and then converted abundances to biovolume using cellular biovolume estimates based on measurements of 20 individuals per species (Table S1) [39]. We assembled three replicate mixtures of control lines and three replicate mixtures of selection lines. From each of the six mixtures, we inoculated two flasks, one with a salt concentration of 0 and one of 6 g NaCl L^−1^. Every four days, we transferred 1 ml of the mixtures into 50 ml of fresh medium, with the flasks continuously shaken at 250 rpm. Over the course of four transfers, we sampled the mixtures at each transfer, fixed the samples with Lugol’s solution, and counted species abundances with an inverted microscope.

We used different approaches to quantify population growth rate in the monoculture assay and population size in the mixture assay (absorbance and abundance counts, respectively) because high-throughput measurement of absorbance with a plate reader allowed us to collect a dense time-series of absorbance data, while microscopical counts were necessary to differentiate species in the mixture assay.

### Adaptation in monoculture vs. adaptation in pairs

To test if competition can influence adaptation to salt stress, we propagated monocultures and pairs of *Anabaena* and *Scenedesmus* with and without increasing salt for 46 transfers (~230 generations), then re-isolated the two species and quantified their competitive ability with and without salt (Fig. 1). We chose the combination of *Anabaena* and *Scenedesmus* because in other species combinations the inferior competitor was outcompeted quickly both with and without increasing salt. In our experimental setup (i.e. a serial transfer experiment with nutrient-replete conditions), competition is mediated by the difference in growth rates, with the species with faster growth rate expected to outcompete the species with slower growth rate. The monocultures and pairs of the two species were part of the selection experiment described above and experienced the same changes in salt concentration except that cultures spent nine more transfers at the final salt concentration (Fig. S2). To quantify how strongly competition affected the abundances of the two species during the selection experiment, we sampled the pairs and monocultures of *Anabaena* and *Scenedesmus* ten times over the course of the selection experiment and microscopically determined the abundances of the two species in Lugol-fixed samples.

After 46 transfers in the selection experiment, we re-isolated *Anabaena* and *Scenedesmus* by plating the cultures on agar and isolating individual colonies. Monocultures were treated the same way as the pairs. Individual colonies were resuspended in liquid Bold’s medium on well plates, and microscopically checked for contamination after a week of growth. We then merged eight clean isolates per sample to one culture. The two isolation steps were conducted in medium without addition of NaCl. We then pre-conditioned the cultures at the two assay environments (0 and 6 g NaCl L^−1^) for two transfer cycles. Hence, before starting the growth and competition assays, the cultures were cultivated under identical environmental conditions for four cycles (two isolation cycles without salt, two transfer cycles in the assay environment) to reduce non-evolutionary effects.

The pre-conditioned cultures were used to start competition assays to test if competition history and salt history influenced competitive ability. To this end, we assembled pairs of *Anabaena* and *Scenedesmus* in the two final salt environments and determined changes in biomass of the two species through time. We conducted two competition assays that differed in how we assembled the pairs (Fig. 1). First, we assembled pairs of co-evolved *Anabaena* and *Scenedesmus*, i.e. lines that had the same competition and salt history. However, because pairs differed in both competitors in this assay (Fig. 1), differences among treatments could not unequivocally be attributed to one of the two species. We thus conducted a second competition assay where we competed each line of *Anabaena* with the same reference line of *Scenedesmus*, and each line of *Scenedesmus* with the same reference line of *Anabaena*. As reference line, we randomly selected one of the three replicate control lines (i.e. competition history: monoculture, salt history: no salt). The second competition assay was started three weeks after the first competition assay; cultures were stored meanwhile as monocultures at their respective salt selection environment and then acclimated for one cycle to the salt assay environment.

In the competition assays, we measured chlorophyll a concentrations of the two competitors as proxy for biomass using a FluoroProbe (bbe Moldaenke, Kiel-Kronshagen, Germany), a fluorometer which can discriminate among algal groups [40]. Before the start of the assay, we calibrated the FluoroProbe with cultures of *Anabaena* and *Scenedesmus* with known chlorophyll a concentrations. We started the competition assay with equal chlorophyll a concentrations (2 μg) of both species in flasks filled with 50 mL of Bold’s medium. We transferred the cultures every 3.5 days for ten transfers in the first competition assay and for four transfers in the second competition assay and measured chlorophyll a concentrations of both species at each transfer.

### Data analysis

All analyses were computed with R version 3.6.2 [41]. We estimated population growth rates (absorbance day^−1^) in monoculture assays by fitting logistic growth curves to absorbance vs. time curves using the R package *grofit* [42]. Growth curves could not be fit when absorbance did not increase; in this case (the three control lines of *Nitzschia* at 6 g NaCl L^−1^) growth rate was set to zero. Some of the growth curves of *Anabaena* were difficult to fit because this species sometimes grew in clumps at high salt, which led to unreliable absorbance measurements. We thus do not present monoculture growth rates of *Anabaena*.

To investigate treatment effects on growth rates, we computed two-way ANOVAs for each of the five species using assay environment and salt history as explanatory variables. It would have been preferential to include ‘line’ as a random effect because each of the six lines (three control lines, three selection lines) was assayed in two environments (with and without salt, respectively). However, computation of linear mixed models with ‘line’ as a random effect often resulted in singular fits because of zero or almost zero variation among lines. We thus dropped the random effect and computed ANOVAs instead. In case of a significant interaction, we additionally computed two separate ANOVAs for each assay environment with salt history as the explanatory variable. We used the function *lm* to compute ANOVAs and the function *Anova* in the *car* package to calculate P-values.

We used the same approach (i.e. ANOVA) to test treatment effects on abundances of species assayed within mixtures, using abundance averaged over the four transfers of the mixture assay as response variable, and assay environment and salt history as explanatory variables. Abundances were ln-transformed prior to analyses, except for abundances of *Pseudokirchneriella* which remained untransformed due to better fit with model assumptions. To test whether competition affected adaptation of *Anabaena* and *Scenedesmus* to salt, we computed three-way ANOVAs for each of the two species and investigated the effects of competition history, salt history and assay environment on relative chlorophyll a (arcsine square-root transformed) in the competition assay. In all our analyses, we did not use model selection to identify important predictor variables, but included all factors manipulated in the experiment as explanatory variables in the model, that is, model structure followed the experimental design.

## Results

### Adaptation of monocultures

After 185 generations of growth with and without increasing salt, respectively, we detected adaptation to salt in two of five species, *Nitzschia* and *Pseudokirchneriella* (Fig. 2, Table S2), that is, lines with a history of salt stress had higher population growth rates at 6 g NaCl L^−1^ than lines without a history of salt stress (ANOVA: *Nitzschia:* F_1,4_= 16.19, P= 0.016; *Pseudokirchneriella:* F_1,4_= 77.70, P= 0.001). In the no-salt assay environment, however, salt history had no effect on the growth rates of these two species (ANOVA: *Nitzschia:* F_1,4_= 0.64, P= 0.467; *Pseudokirchneriella:* F_1,4_= 2.78, P= 0.171). Among the remaining three species, *Scenedesmus* and *Navicula* had reduced growth rates in the high-salt assay environment irrespective of salt history, while *Synechococcus* was unaffected by both salt history and assay environment (Fig. 2, Table S2).

**Figure 2:**
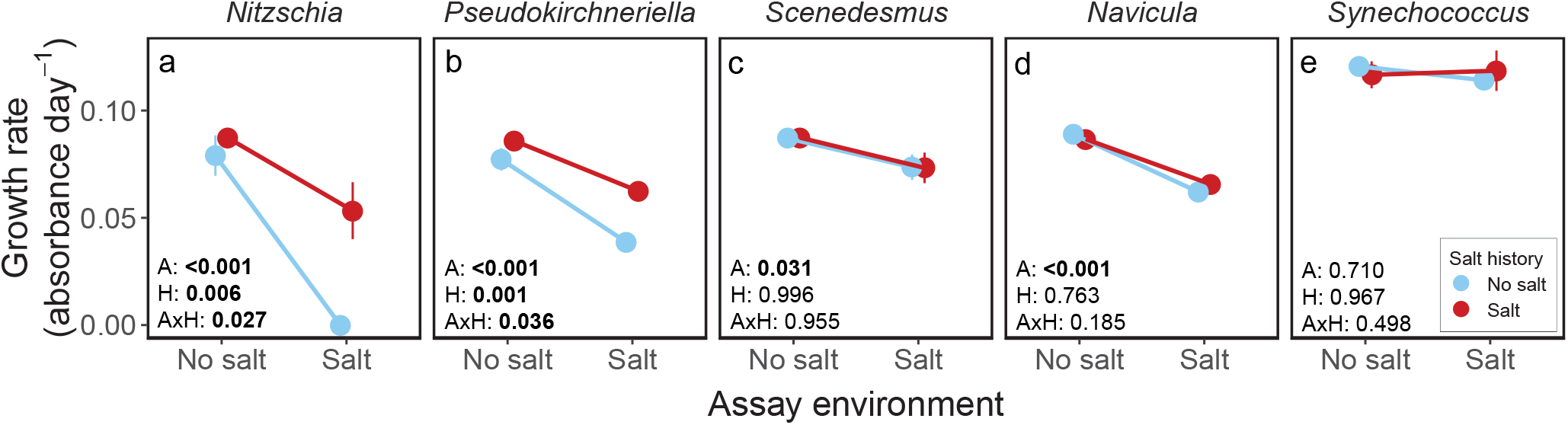
Effects of salt history and assay environment on the monoculture growth rates of five species of microalgae (sorted by salt tolerance). Control lines (blue) have a salt history of 37 transfers at 0 g NaCl L^−1^, selection lines (red) have a salt history of 37 transfers at increasing salt stress with a final concentration of 6 g NaCl L^−1^. Salt concentrations in the two assay environments were the same as in the final selection environments, i.e. 0 and 6 g NaCl L^−1^, respectively. Inserts give the P-values for assay environment (A), salt history (H), and their interaction (AxH). Bold font denotes P-values < 0.05. For F-statistics see Table S2. Values are means ± SE, n = 3.

### Mixture assay

When assaying mixtures of either control or selection lines, we found that the effect of salt history varied among species and depended on assay environment (Fig. 3 a-c, Table S3). Most species declined in abundance over the four transfers of the assay, except for *Synechococcus*, which increased (Fig. S3). Salt history affected the abundance of three species: *Nitzschia, Pseudokirchneriella, Scenedesmus* (Fig. 3 a-c, Table S3). In *Nitzschia*, lines with a history of salt stress had lower average abundance than control lines when assayed without salt (58% lower abundance; F_1,4_= 25.97, P= 0.007). At high salt, however, the effect of salt history on average abundance of *Nitzschia* was not significant (F_1,4_= 4.54, P= 0.10) and control and selection lines declined similarly fast (Fig. S3a). In *Pseudokirchneriella*, selection lines had lower average abundance than control lines in the no-salt assay environment (24% lower abundance; F_1,4_= 24.11, P = 0.008), but 2.5 times higher abundance than control lines in the high-salt assay environment (F_1,4_= 14.90, P = 0.018). However, these treatment effects on *Pseudokirchneriella* did not persist until the end of the mixture assay: within four transfers, *Pseudokirchneriella* went extinct or declined to low abundance irrespective of salt history and assay environment (Fig. S3b). In *Scenedesmus*, a history of salt stress had no effect on average abundance in the no-salt assay environment (F_1,4_= 1.73, P = 0.258), but a positive effect at high salt (1.7 times higher abundance; F_1,4_= 11.86, P = 0.026). Among the remaining three species, abundance of *Anabaena* was negatively affected by salt irrespective of salt history (Fig. S3c, Table S3), while *Navicula* and *Synechococcus* were unaffected by both salt history and assay environment (Fig. 3d, e, Table S3).

**Figure 3:**
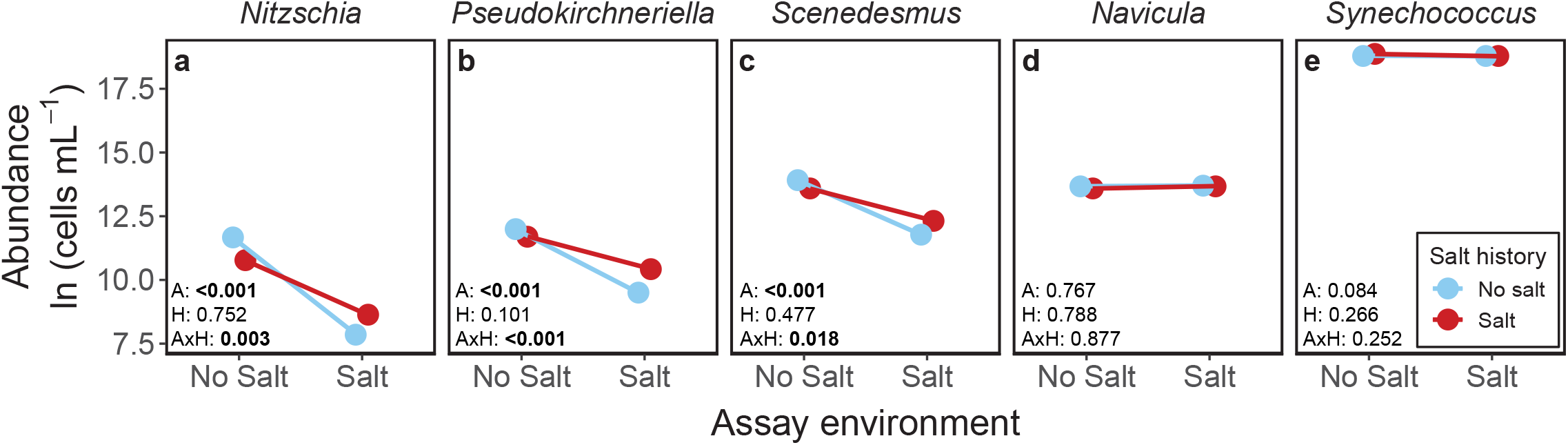
Effects of salt history and assay environment on species abundances in the mixture assay. Mixtures of six species were assembled either from control lines (blue) or from selection lines (red). Species abundances were averaged over four transfers in assay environments with and without salt, respectively. Inserts give the P-values for assay environment (A), salt history (H), and their interaction (AxH). For F-statistics see Table S3. Abundances were ln-transformed prior to analyses except for *Pseudokirchneriella*. For consistency with Fig. 2, we do not show *Anabaena*, which was also part of the communities. For time-resolved data of all six species, see supplementary figure S4. Values are means ± SE (SE too small to be visible), n = 3.

### Adaptation in monoculture vs. adaptation in pairs

Interspecific competition strongly influenced species abundances during the selection experiment, but did not lead to any evolutionary responses. In the selection experiment, *Anabaena* reduced the abundance of *Scenedesmus* as long as salt concentration was low (Fig. S4b). However, at the final salt concentration of 6 g NaCl L^−1^ the competitive hierarchy reversed and *Anabaena* was negatively affected by *Scenedesmus* (Fig. S4a). This change in competitive hierarchy in response to salt stress was also apparent in the competition assay: *Anabaena* dominated without salt (Fig. 4a), while *Scenedesmus* dominated at high salt (Fig. 4b), irrespective of competition history.

**Figure 4:**
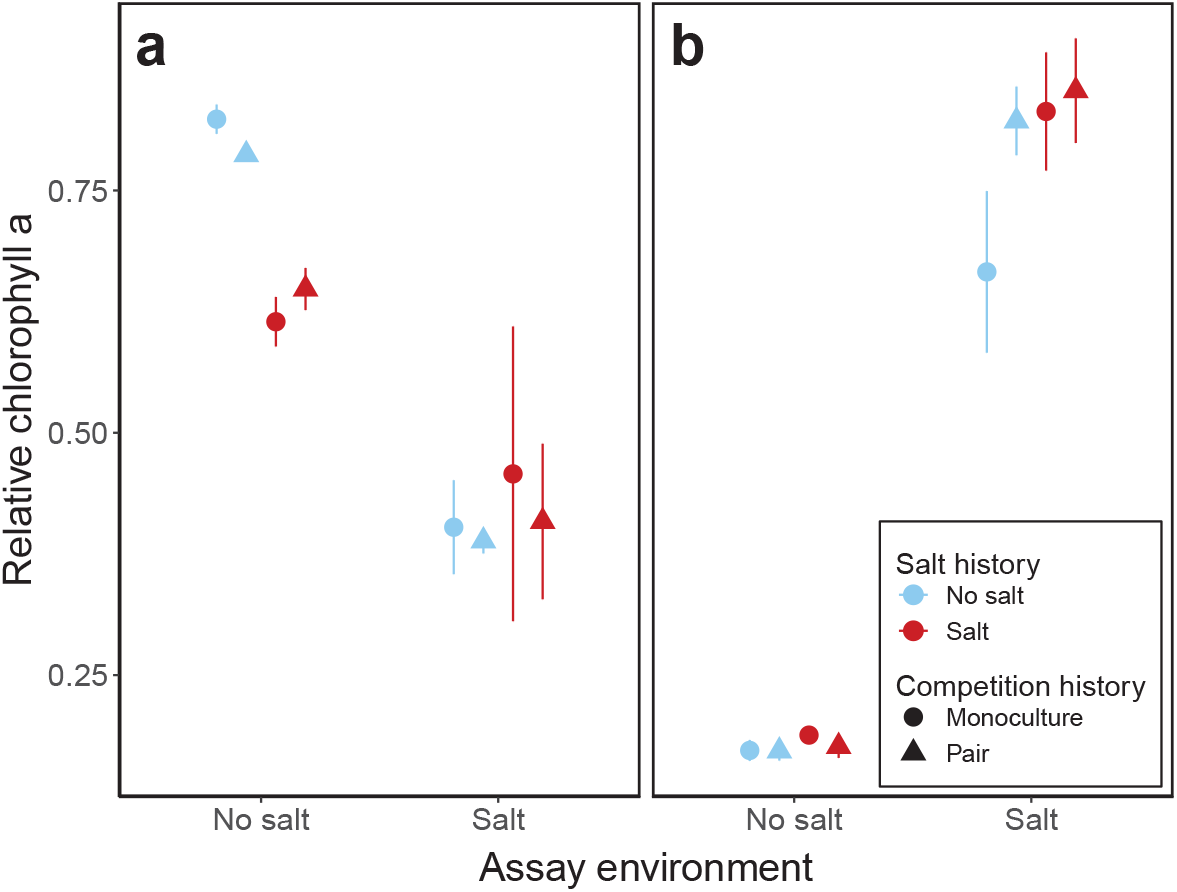
Effects of competition history, salt history, and assay environment on competitive abilities of *Anabaena* and *Scenedesmus*. Relative chlorophyll a of (a) *Anabaena* in competition with a reference line of *Scenedesmus*, and (b) of *Scenedesmus* in competition with a reference line of *Anabaena*. Values are means ± SE, n = 3.

The two competition assays revealed that lines of *Anabaena* with a history of salt stress had reduced competitive ability in the no-salt assay environment. When assayed with co-evolved competitors, high-salt lines of *Anabaena* had lower relative chlorophyll a than no-salt lines in the no-salt assay environment, and consequently high-salt lines of *Scenedesmus* had higher relative chlorophyll a than no-salt lines (salt history × assay: F_1,16_ = 13.47, P = 0.002; Table S4, Fig. S5). Because both competitors differed among treatment combinations, this experiment did not allow detecting if the observed pattern was due to altered competitive ability of *Anabaena* or of *Scenedesmus*. However, when assayed in competition with a reference line, only *Anabaena* was affected by salt history, again with lower relative chlorophyll a of high-salt than no-salt lines in the no-salt assay environment (salt history × assay: F_1,16_ = 5.78, P = 0.029, Table S5, Fig. 4a). As a consequence, in the no-salt assay environment the biomass (i.e. chlorophyll a concentration) of the competing reference line varied by a factor of 1.9 depending on the evolutionary history of *Anabaena* (Fig. S6a).

## Discussion

With our experiments we addressed a range of scenarios that can arise from the interplay of competition and adaptation to an abiotic stressor. In the first experiment, we found that competition can alter how adaptation to an abiotic stressor influences population dynamics (Fig. 2, 3): effects of salt history on population growth rate in monoculture did not necessarily lead to increased population size within communities. Conversely, some evolutionary effects, specifically, reduced population size of selection lines without salt, appeared only within mixtures of species, not in monoculture. In the second experiment, we found no effect of competition on adaptation to salt stress (Fig. 4). However, salt history influenced competition: after selection at increasing salt, one of two species had reduced competitive ability without salt (Fig. 4a). Collectively, our results highlight that ecological and evolutionary processes interactively shape communities in changing environments, but that these eco-evolutionary effects can be diverse and species-specific.

Species varied in their evolutionary response to salt stress. The two species with the lowest limit of salt tolerance adapted to salt (Fig. 2a, b), while species that were less negatively affected by salt showed no sign of adaptation (Fig. 2 c-e). This finding is in line with the prediction that species with narrow tolerance of a stressor should evolve more than species with broader tolerance because they experience stronger selection pressure [43]. Species-specific differences in genetic variance and covariance could be additional reasons for variation among species in the rate of evolution [43, 44]. Accordingly, an experiment that investigated thermal evolution in three phytoplankton species found largest fitness improvements in the species with highest population density and the least complex genome [45]. Given that only two species adapted in our experiment, our sample size is not large enough to evaluate if species-specific differences other than ecological tolerance influenced the evolutionary response to salt. Such variation among species in the ability to adapt to environmental change could add further complexity to adaptation in a community context.

Our experiments suggest that competition can modulate how adaptation to abiotic change influences population dynamics. When assayed at high salt in monoculture, two species (*Nitzschia* and *Pseudokirchneriella*) showed signatures of adaptation, that is, higher population growth rate of selection lines than control lines (Fig. 2a, b). When assayed at high salt within mixtures, however, only one of these two species (*Pseudokirchneriella*) had higher population size of selection than control lines (Fig. 3b), and this positive effect of salt history was only transient (Fig. S3b). In contrast, control and selection lines of *Nitzschia* had similar population size and declined equally fast within salt-stressed mixtures (Fig. 3a, Fig. S3a). Similarly, adaptation of a marine alga to acidification was not associated with increased population size in acidified communities [20]. Such a pattern can emerge when competition changes the fitness landscape [18]; thereby adaptation of a species to abiotic change improves its fitness only in the same community context in which it has evolved [18]. In our mixtures, low competitive ability of the two species that adapted to salt might have also contributed to this pattern. *Nitzschia* and *Pseudokirchneriella* were the weakest competitors in the community and declined to extinction or to low abundance irrespective of salt history and salt environment (Fig. S3). It remains to be determined if effects of adaptation on population dynamics are less contingent on community context in stronger competitors. In addition to low competitive ability, low starting abundance could have contributed to the fast extinction of *Nitzschia:* this species was the largest in our community (Table S1) and was thus inoculated with the lowest abundance among the six species. Taken together, the positive effect of adaptation to abiotic change on population size may be reduced when community context changes; such a scenario could result from immigration of novel competitors in response to environmental change [27, 28, 46].

We also found evolutionary effects that appeared only within mixtures. In *Nitzschia* and *Pseudokirchneriella*, adaptation to salt was associated with costs (i.e. reduced population size) in the no-salt environment when these species were assayed within mixtures (Fig. 3a, b), but not when assayed in isolation (Fig. 2a, b). A possible mechanism underlying such a pattern is a trade-off between traits that mediate stress tolerance and competition [34]. We did not measure traits and can thus only speculate about potential mechanisms. Tolerance of high salt concentrations is conferred by the ability to efficiently export ions (Na^+^ and Cl^−^) and accumulate compatible solutes, i.e. organic molecules that maintain osmotic balance without interfering with metabolism [47]. High energetic costs for ion export and for synthesis of compatible solutes [48] may result in reduced competitive ability. However, when adaptation to an abiotic stressor occurs within communities, rather than in isolation as in our experiment, reduced competitive ability might not evolve. Our experiment thus simulates a scenario in which environmental change is followed by a change in community context, which could result from immigration of novel competitors or from changes in local community composition due to species-specific differences in environmental tolerance. To elucidate if adaptation to an abiotic stressor is associated with changes in traits that mediate biotic interactions, a trait-based approach would be an interesting avenue for future experiments.

Competition had no effect on adaptation to salt stress (Fig. 4). We had predicted that competition during the selection experiment would constrain adaptation to salt because of reduced population size and/or increased genetic constraints [23, 26]. Alternatively, competition could facilitate adaptation to abiotic stress by increasing the strength of selection [25]. The two species that we used to test this question, *Anabaena* and *Scenedesmus*, adapted to salt neither in monoculture nor in pairs. Similarly, an experiment that manipulated CO_2_ enrichment and competition in a selection experiment with two algae species found neither adaptation to CO_2_ nor to competition [19]. In our experiment, low selection pressure could explain why the monocultures did not adapt to salt, given that the two species had not reached their limit of salt tolerance in the selection experiment. However, the combination of salt and competition in the selection experiment drove the inferior competitor to the brink of extinction (Fig. S4), suggesting that the inferior competitor experienced strong selection pressure to evolve faster growth rates. Yet, we did not find any effect of competition history on adaptation. One possible explanation is that the competitive hierarchy in flasks with increasing salt changed shortly before the end of the selection experiment when the final salt concentration had been reached, from dominance of *Anabaena* to dominance of *Scenedesmus*. Maybe relaxed selection pressure on *Scenedesmus* towards the end of the experiment and only a short phase of strong selection on *Anabaena* explain why competition had no effect on the evolutionary response of these two species.

Evolutionary change in response to salt influenced competitive ability in one of two species (Fig. 4). We had predicted that a history of salt stress would lead to increased competitive ability at high salt. In contrast, however, strains of *Anabaena* with a history of salt stress had reduced competitive ability in the no-salt assay environment. Such a pattern is an indication of conditionally deleterious mutations [49], i.e. mutations that are neutral in the local environment and deleterious in an alternative environment. Because we have neither molecular nor trait data, we cannot elucidate potential mechanisms behind the observed mal-adaptation in the ancestral environment. The evolutionary response of *Anabaena* was independent of competition history and of whether fitness was assayed in competition with a co-evolved or a naïve competitor (Fig. 4, Fig. S5). The ecological effect of this evolutionary response to salt is also illustrated by the response of the competing reference line: in the no-salt assay, the chlorophyll a concentration attributable to the reference line of *Scenedesmus* was 1.9 times higher when competing with selection lines of *Anabaena* than when competing with control lines (Fig. S6a). Overall, our results suggest that evolutionary responses to abiotic change can influence biotic interactions, such that populations with different histories of abiotic stress might vary in their effects on communities and ecosystems.

The emerging field of eco-evolutionary dynamics investigates the effects of evolution on populations, communities, and ecosystems [6]. Research on adaptation to abiotic change usually explores the effect of evolution on population dynamics [4, 7, 11], while studies that investigate the effect of evolution on species interactions often focus on biotic selective agents [50–52]. Here we showed that evolutionary responses to an abiotic selective agent can alter biotic interactions (Fig. 4a, Fig. S6a). Moreover, we found that competition can modulate the effect of adaptation on population dynamics (Fig. 2, 3), suggesting that adaptation to abiotic change may not enhance performance of a species when community context changes [18] e.g. because of immigrating competitors [28]. We have observed a range of complexities arising from adaptation in a multi-species context despite using a fairly simple model system. Higher diversity and trophic structure of natural systems will pose an even greater challenge to understanding eco-evolutionary dynamics in changing environments [33, 53]. Environmental fluctuation is another complexity of natural systems that we were not able to address in our experiment. Temporary environmental amelioration could promote adaptation to abiotic change by increasing population size and thus the supply of beneficial mutations [11], but could also reduce the potential for adaptation by relaxing selection pressure [54]. Taking these complexities into account will be crucial in future work to further increase our understanding of adaptation to abiotic change in a community context.

## Acknowledgements

We thank Etienne Low-Décarie and Graham Bell for input on the design and analysis of the experiment, and Yinci Yan for help with the implementation of the experiment.

## Funding

This work was supported by an Erwin-Schrödinger scholarship of the Austrian Science Fund (FWF), grant no. J3265-B16 to R.L., and an NSERC Discovery Grant to G.F.F.

## Notes

### Competing Interest Statement

The authors have declared no competing interest.

