## Supplementary tables and figures for "Adaptation and competition in deteriorating environments"

### Supplemental tables and figures

**Table S1:** Cellular biovolume of the six species of microalgae used in the experiments.

| Species | Biovolume ($\mu m^3$) |
|---|---|
| *Anabaena variabilis* | 12 |
| *Navicula pelliculosa* | 28 |
| *Nitzschia palea* | 112 |
| *Pseudokirchneriella subcapitata* | 38 |
| *Scenedesmus acutus* | 57 |
| *Synechococcus leopoliensis* | 5 |

**Table S2:** Results of two-way ANOVAs testing the effects of assay environment and salt history on monoculture growth rates of five species of microalgae. P < 0.05 in bold.

| | Assay | | History | | Assay × History | |
|---|---|---|---|---|---|---|
| Species | $F_{1,8}$ | *P* | $F_{1,8}$ | *P* | $F_{1,8}$ | *P* |
| *Nitzschia* | 45.51 | **<0.001** | 13.52 | **0.006** | 7.27 | **0.027** |
| *Pseudokirchneriella* | 110.43 | **<0.001** | 30.13 | **0.001** | 6.35 | **0.036** |
| *Scenedesmus* | 6.78 | **0.031** | 0.00 | 0.996 | 0.00 | 0.955 |
| *Navicula* | 142.69 | **<0.001** | 0.10 | 0.763 | 2.11 | 0.185 |
| *Synechococcus* | 0.15 | 0.710 | 0.00 | 0.967 | 0.50 | 0.498 |

**Table S3:** Results of ANOVAs testing for effects of assay environment and salt history on species abundances in the mixture assay. Abundances were averaged over four transfers and ln-transformed prior to analyses, except for abundances of *Pseudokirchneriella* which remained untransformed due to better fit with model assumptions. P < 0.05 in bold.

| | Assay | | History | | Assay × History | |
|---|---|---|---|---|---|---|
| Species | $F_{1,8}$ | *P* | $F_{1,8}$ | *P* | $F_{1,8}$ | *P* |
| *Nitzschia* | 223.72 | **<0.001** | 0.11 | 0.752 | 17.27 | **0.003** |
| *Pseudokirchneriella* | 596.28 | **<0.001** | 3.43 | 0.101 | 38.85 | **<0.001** |
| *Anabaena* | 58.13 | **<0.001** | 1.65 | 0.235 | 1.22 | 0.301 |
| *Scenedesmus* | 131.29 | **<0.001** | 0.56 | 0.477 | 8.79 | **0.018** |
| *Navicula* | 0.09 | 0.767 | 0.08 | 0.788 | 0.03 | 0.877 |
| *Synechococcus* | 3.89 | 0.084 | 1.43 | 0.266 | 1.53 | 0.252 |

**Table S4:** Results of three-way ANOVA testing for effects of assay environment, salt history, and competition history on relative chlorophyll a of *Anabaena* and *Scenedesmus* in competition with co-evolved *Scenedesmus* and *Anabaena*, respectively. Relative chlorophyll a was averaged over transfers 4-10. The data was arcsin square root transformed prior to analysis. P-values < 0.05 in bold.

| | $F_{1,16}$ | P-value |
|---|---|---|
| Assay environment | 247.185 | **< 0.001** |
| Salt history | 42.806 | **< 0.001** |
| Competition history | 1.875 | 0.190 |
| Assay environment × Salt history | 13.496 | **0.002** |
| Assay environment × Competition history | 1.856 | 0.192 |
| Salt history × Competition history | 0.576 | 0.459 |
| Assay environment × Salt history × Competition history | 2.754 | 0.117 |

**Table S5:** Results of three-way ANOVAs testing the effects of assay environment, salt history, and competition history on relative chlorophyll a of *Anabaena* and *Scenedesmus* in competition with a reference line of *Scenedesmus* and *Anabaena*, respectively. The data was arcsin square root transformed prior to analysis. P-values < 0.05 in bold.

| | *Anabaena* | | *Scenedesmus* | |
|---|---|---|---|---|
| | $F_{1,16}$ | P | $F_{1,16}$ | P |
| Assay environment | 44.342 | **< 0.001** | 284.389 | **< 0.001** |
| Salt history | 2.801 | 0.114 | 3.036 | 0.101 |
| Competition history | 0.141 | 0.713 | 1.368 | 0.259 |
| Assay environment × Salt history | 5.784 | **0.029** | 1.983 | 0.178 |
| Assay environment × Competition history | 0.063 | 0.805 | 1.885 | 0.189 |
| Salt history × Competition history | 0.065 | 0.802 | 1.062 | 0.318 |
| Assay environment × Salt history × Competition history | 0.357 | 0.559 | 0.714 | 0.411 |

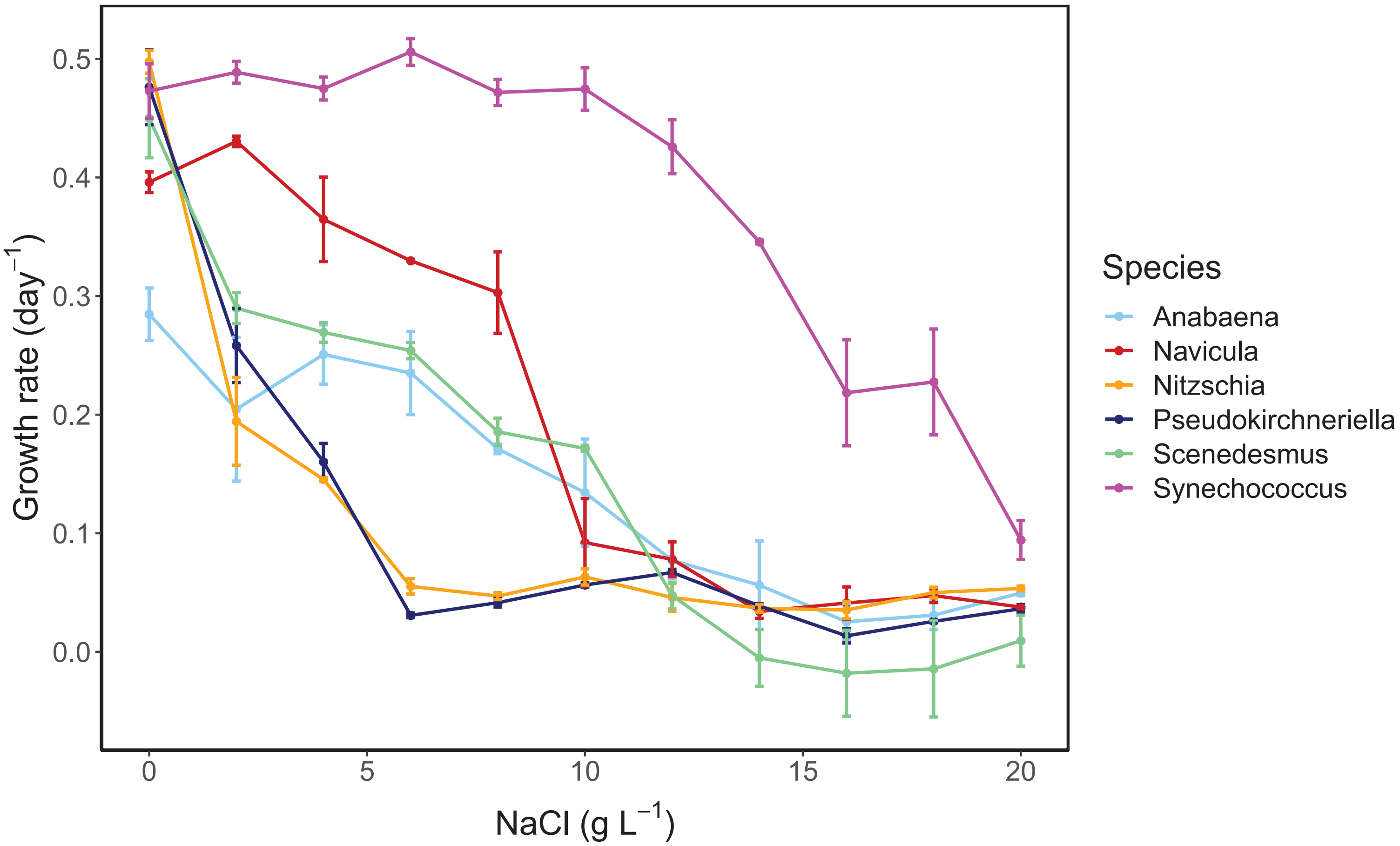


**Fig. S1:** Estimates of salt tolerance. We measured growth rates of ancestral lines along a salt gradient from 0 to 20 g NaCl $L^{-1}$ in 2 g increments. Because the salt tolerance experiment was conducted in different containers (48-well plates) than the selection experiment (flasks), the salt tolerance data are only coarse estimates of salt tolerance in the main experiment. Later assays in flasks indicated that cultures often grew better in flasks; while both flasks and well plates were continuously shaken, shaking of well plates did not result in complete homogenization of cultures, which may have led to lower nutrient supply compared to flasks. Moreover, due to evaporation and a greater risk of cross-contamination of cultures on well plates, the salt tolerance experiment had to be terminated after seven days, while the assay in flasks could be maintained for several weeks. Consequently, when species had long lag phases (control lines of *Pseudokirchneriella* at 6 g NaCl $L^{-1}$), their ability to grow was captured in the growth assay of the main experiment (Fig. 2b), but not in the salt tolerance experiment. Due to the shorter duration of the salt tolerance experiment, cultures often did not reach carrying capacity, which is why we could not fit logistic growth curves to this dataset. Instead, we estimated growth rates by fitting linear models to the exponential part of the absorbance vs. time curves, using ln-transformed absorbance data as response variable. This graph thus shows the per capita growth rate ($day^{-1}$), while the results of the growth assay presented in the main text (Fig. 2) are displayed as population growth rate (absorbance $day^{-1}$). Values are means ± SE, n=2.

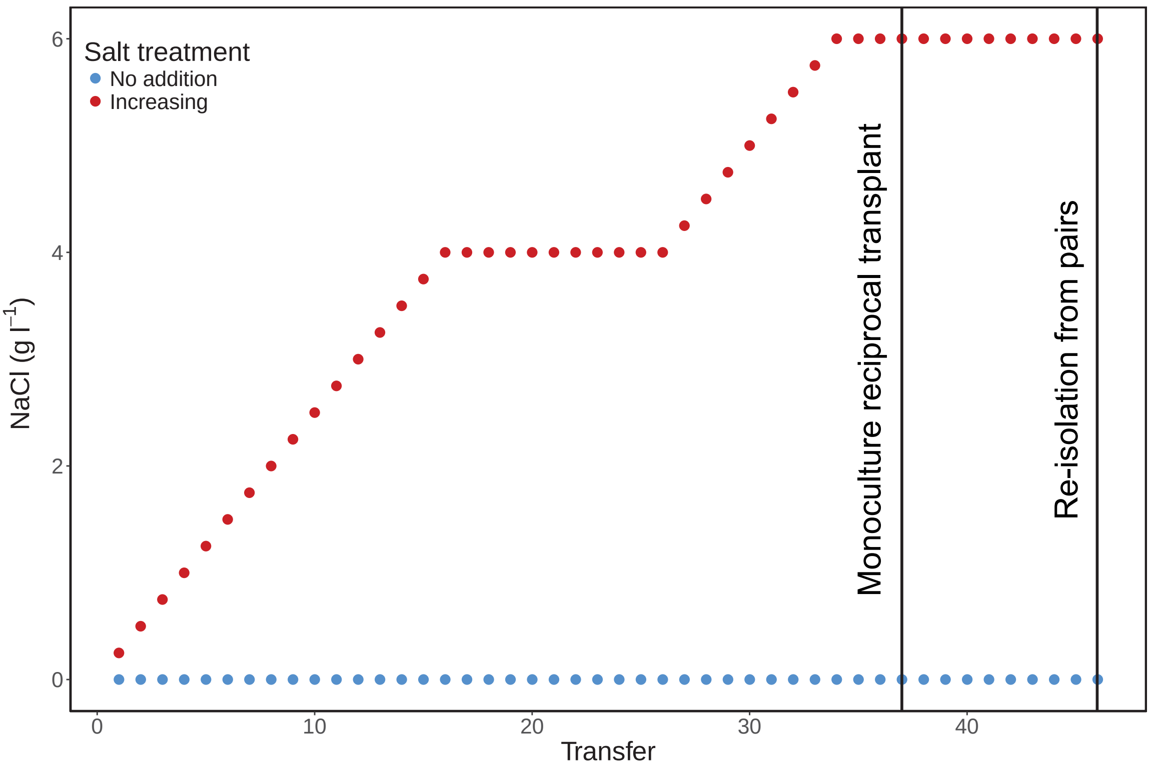


**Fig. S2:** Salt concentration over the course of the selection experiment. No salt was added to the control lines (blue), while the selection lines were exposed to increasing concentrations of NaCl (red). At transfer 37, the cultures were subsampled to start the acclimation phase for the monoculture and community assays. At transfer 46, monocultures and pairs of *Anabaena* and *Scenedesmus* were plated on agar for isolation and subsequent assays.

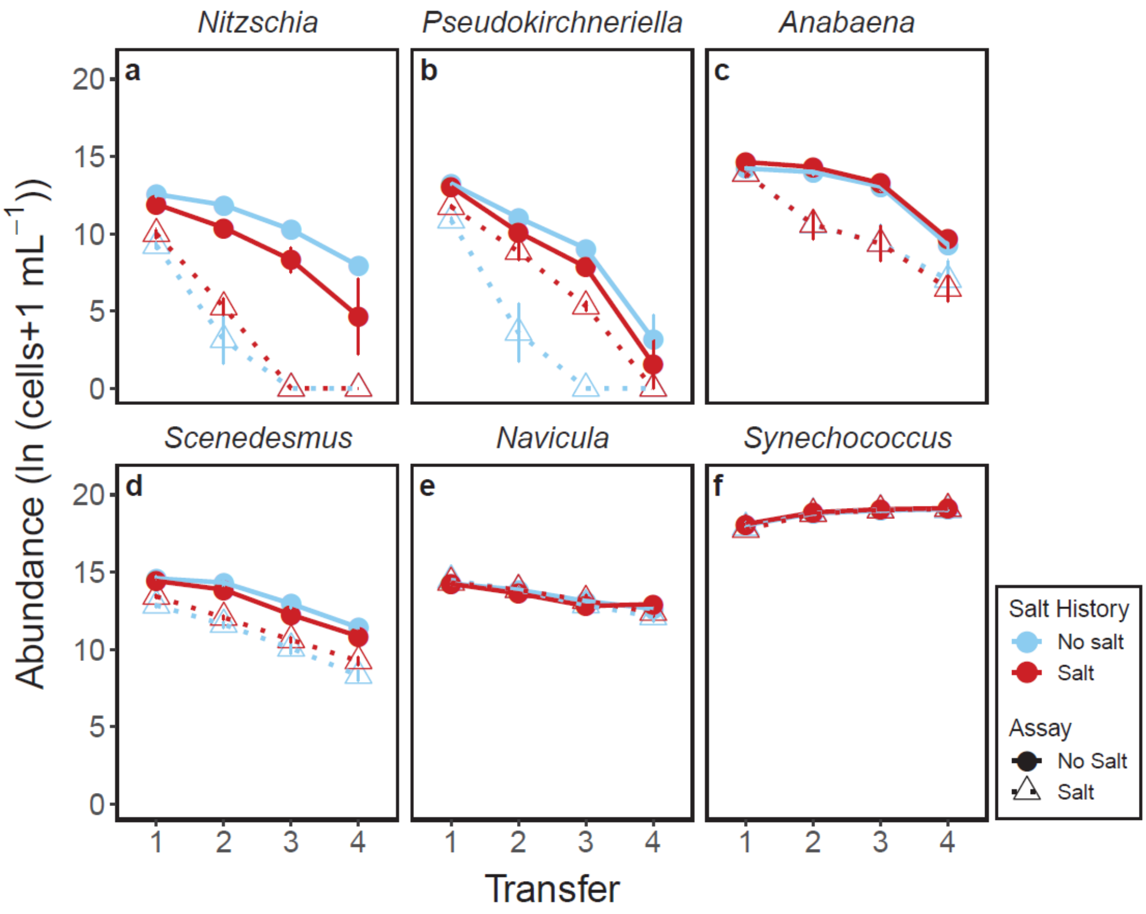


**Fig. S3:** Effects of salt history and assay environment on species abundances in the mixture assay. Mixtures of six species were assembled either from control lines (blue) or from selection lines (red). Species abundances were quantified over four transfers in assay environments without salt (solid circles, solid lines) and with salt (open triangles, dotted lines), respectively. Species are sorted based on salt tolerance, from species with low salt tolerance (*Nitzschia*, *Pseudo-kirchneriella*), to medium salt tolerance (*Anabaena*, *Scenedesmus*, *Navicula*), to high salt tolerance (*Synechococcus*). Values are means ± SE, n = 3.

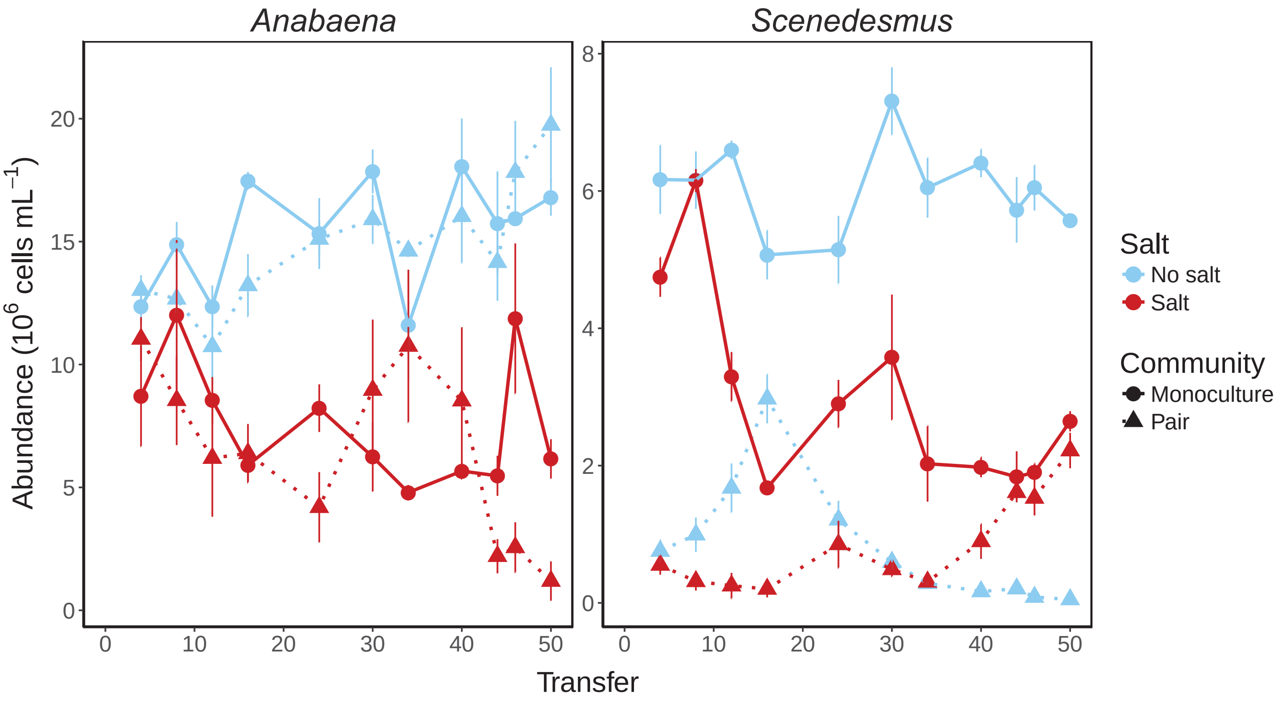


**Fig. S4:** Abundances of *Anabaena* and *Scenedesmus* over the course of the selection experiment. The two species were propagated in monocultures and in pairs, in medium with or without increasing salt concentration. Values are means ± SE, n=3.

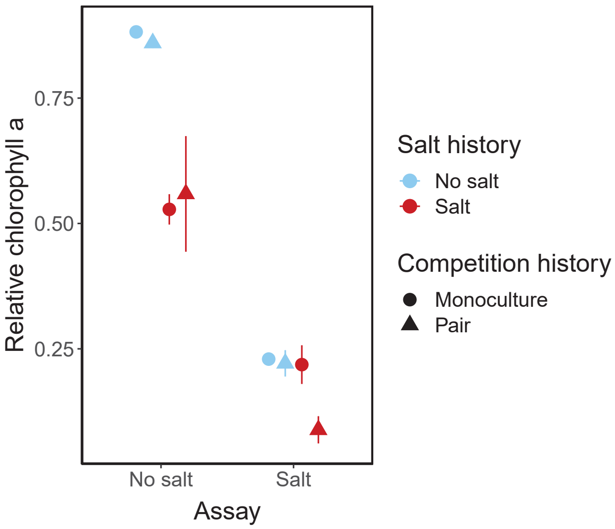


**Fig. S5:** Effects of competition history, salt history, and assay environment on relative chlorophyll a of *Anabaena* in competition with co-evolved *Scenedesmus* (i.e. *Scenedesmus* with the same salt and competition history). Competitive ability was estimated as relative chlorophyll a averaged over transfers four to ten of the competition assay. Values are means ± SE, n = 3.

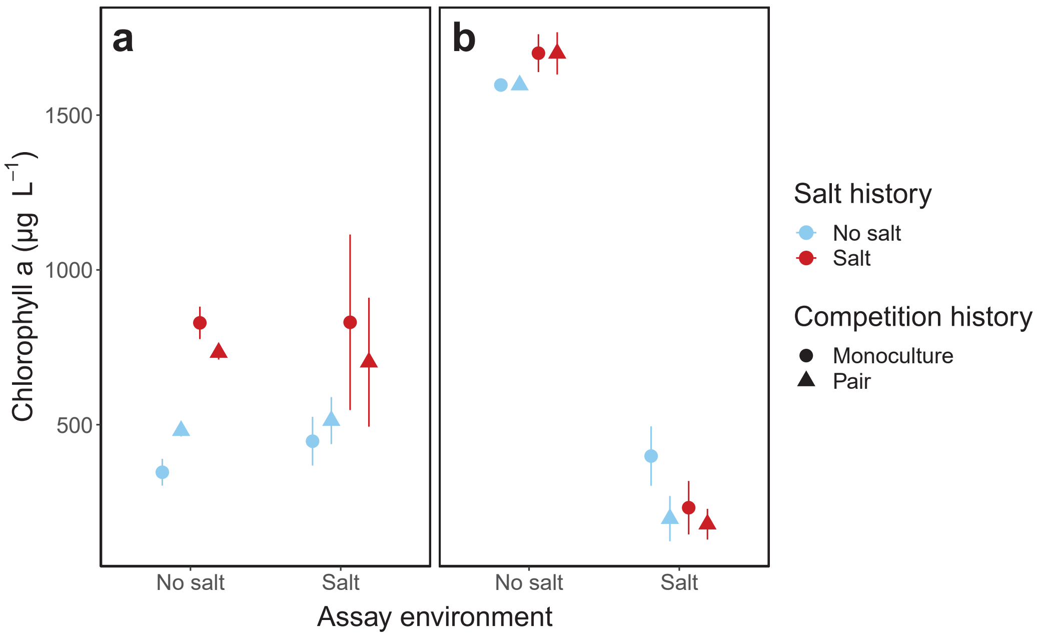


**Fig. S6:** Chlorophyll a concentration of (a) the reference line of *Scenedesmus*, and (b) the reference line of *Anabaena*, as a function of the evolutionary history of its competitor. Values are means ± SE, n = 3.
